# RdDM pathway is required for *Tobamovirus*-induced symptomatology production

**DOI:** 10.1101/2020.01.21.912923

**Authors:** Melisa Leone, Diego Zavallo, Andrea Venturuzzi, Sebastián Asurmendi

## Abstract

Small RNAs (sRNA) are important molecules for gene regulation in plants and play an essential role in plant-pathogen interactions. Researchers have evaluated the relationship between viral infections as well as the endogenous accumulation of sRNAs and the transcriptional changes associated with the production of symptoms, little is known about a possible direct role of epigenetics, mediated by 24-nt sRNAs, in the induction of these symptoms.
With the use of different RNA directed DNA methylation pathway mutants and triple demethylase mutants, here we demonstrate that the disruption of RdDM pathway during viral infection produced alterations in the plant transcriptomic changes (because of the infection) and in symptomatology.
This study represents the initial step in exposing that DNA methylation directed by endogenous sRNAs has an important role, uncoupled to defense, in the production of symptoms associated with plant-virus interactions.

**Significance statement:** The crop yield losses induced by phytoviruses are mainly associated with the symptoms of the disease. DNA modifications as methylation, can modulate the information coded by the sequence, process named epigenetics. Viral infection can change the expression patterns of different genes linked to defenses and symptoms. This work represents the initial step to expose the role of epigenetic process, in the production of symptoms associated with plants-virus interactions.

## Introduction

Diseases associated with viruses produce diverse symptoms and numerous losses in crop losses. Phytoviruses induce different metabolic and physiological alterations that in many cases result in symptomatic disease phenotypes. Viral infections can cause growth retardation, leaf mosaic, yellowing, necrosis and various developmental disorders (Wang *et al*., 2012). For this reason, understanding the molecular bases associated to symptomatology production is essential to generate new biotechnological strategies in search of resistant plants (Nicaise, 2014).

The *Tobamoviruses* genus includes 37 species of viruses known to infect many crops, such as potatoes, tomatoes, cucumbers, tobacco, among others. Tobacco mosaic virus strain Cg (TMV-Cg) and Oilseed rape mosaic virus (ORMV) are two strains (with more than 90% genome sequence homology) of the *Tobamovirus* genus that infect *Arabidopsis thaliana* plants (Aguilar *et al*., 1996). Briefly, during infection, the virus first entries the organism after mechanically damaging the cell wall and the plasma membrane. Soon after, Capside Protein (CP) begins to be disengaged from the virion and, then, the first protein translated directly from de genome, the replicase, initiates the replication of the viral genome (Shaw, 1999). The movement of the virus depends on viral Movement Proteins (MP), although other viral proteins are responsible for this process in some other species.

An efficient viral infection requires the interaction between viral proteins and host cell factors that causes a manipulation of metabolic pathways and coordinates the biochemical interactions promoting infection. The recognition of viral double stranded RNA (dsRNA) molecules by the plant defense system leads to the induction of RNA silencing (Niehl *et al*., 2016; Niehl and Heinlein, 2019). However, many plant viruses evade this silencing process and induce viral counter defense (Vance and Vaucheret, 2001; G. Conti *et al*., 2017). Finally, as a result of the mentioned viral-plant interaction, systemic symptoms appear. The causes of symptom induction are diverse and, moreover, many depend on the interference with the normal physiological processes of plants occurring upon defense response (Pallas and García, 2011).

In addition, viral infection is associated with global transcriptome reprogramming and changes in hormonal pathways as well as with the increase of the accumulation of metabolites and antioxidant compounds that produce disease phenotypes (Westwood *et al*., 2013; Dastogeer *et al*., 2018; Bazzini *et al*., 2011). For instance, chlorosis is a characteristic symptom widely correlated with modifications induced by viruses as a result of changes in the quantity, size or structural alterations of chloroplasts and with the global repression of photosynthetic genes (Bilgin *et al*., 2010; Li *et al*., 2016; Bhattacharyya and Chakraborty, 2018).

RNA silencing is a transcriptome regulation mechanism that uses small RNAs (sRNAs) as guides and that can act transcriptionally by cytosine DNA methylation via Transcriptional Gene Silencing (TGS) or post-transcriptionally (PTGS) by cleavage of RNA targets (Borges and Martienssen, 2015; Castel and Martienssen, 2013; Kørner *et al*., 2018) to regulate gene expression, modify chromatin topology or defend against viral infections (Baulcombe, 2004). In this context, the RNA-directed DNA methylation (RdDM) pathway is a methylation mechanism that can occur in different cytosine sequence contexts directed by 24-nt sRNAs (Gallego-Bartolomé *et al*., 2019; Law and Jacobsen, 2010). The canonical RdDM pathway requires the coordination of two RNA polymerases (Pol IV-Pol V). Pol IV RNA transcripts generated dsRNAs and RNA-dependent RNA polymerase 2 (RDR2) forms the dsDNAs that are processed by Dicer like 3 (DCL3) to produce 24-nt sRNAs. Then, 24-nt sRNAs are loaded into argonaute 4 (AGO4) and this combined structure interacts with Pol V to, finally, lead the methylation of DNA target sequences through the interaction with a methyltransferase (Xie and Yu, 2015; He *et al*., 2014).

DNA methylation is an epigenetic phenomenon that plays a key regulatory role in many biological processes, including transposon silencing and responses to biotic and abiotic stresses. Usually, methylation of promoters, transposons or repeat sequences is linked to TGS effects, whereas gene-body methylation in some cases impacts on transcription, and, although still unclear, it seems to be positively correlated with active transcription (Liang *et al*., 2019; To *et al*., 2015). The insertion of transposons near or into genes can alter gene expression, which in some cases can produces phenotypic consequences (Bewick and Schmitz, 2017). For example, a successful heat response in *Arabidopsis* depends on the integrity of epigenetic pathways and provides evidence that heat-dependent gene expression is influenced by transposon sequences located near genes linked to heat stress (Popova *et al*., 2013). Achour et al. (2019) showed that prolonged low temperature exposure promotes the activation of genes involved in the regulation of DNA methylation. Thus, this transcriptional activation together with the extensive change of methylation in transposons along the genome shows the importance of hypermethylation of TEs in response to abiotic stress.

In addition, DNA methylation also plays an important role in defense response. In recent years, several studies have recognized the link between epigenetics and immunity. For instance, defective Pol V mutants show greater resistance to disease against bacterial pathogen *Pseudomonas syringae* (López *et al*., 2011). In other study, demethylation activated Xa21, a gene that confers resistance to bacterium Xanthomonas *oryzae pv.* in rice and, furthermore, the progeny inherited both hypomethylation and the acquired phenotype (Akimoto *et al*., 2007). In addition, methylation of pericentromeric TEs could affect the expression of PRR/NLR genes (Cambiagno *et al*., 2018). In recent years, researchers have published several studies about the effects of DNA methylation on antiviral defense. The infection by TRV and Cucumber Mottled Green Mosaic Virus (CGMMV) in Arabidopsis and watermelon, respectively, produces a deregulation of key components of the RdDM machinery, thus evidencing an important role of methylation in antiviral defense (Diezma-Navas *et al*., 2019; Sun *et al*., 2019). Another study revealed a novel regulatory mechanism, whereby Cotton Leaf Curl Multan virus (CLCuMuV) C4 protein suppresses TGS and PTGS mediated by SAMS (a central enzyme in the methyl cycle), which leads to an improved viral infection in the plant (Ismayil *et al*., 2018). Although the molecular mechanisms involved in the production of symptoms because of viral induction have been widely studied, they are still poorly understood.

In this study, we assessed the role of epigenetics in the production of symptoms in *Arabidopsis thaliana* inoculated with two *Tobamoviruses* strains (ORMV AND TMV-Cg). We detected that viral infection modified the expression of genes mapped by differentially accumulated (DA) 24-nt sRNAs. The impact of this alteration is not directly linked to plant defense, although it affects the emergence of symptoms in an RdDM-dependent manner. Our findings reveal a role of DNA methylation in viral symptomatology.

## Results

Because 24-nt sRNAs play an important role in plant immunity through Transcriptional Gene Silencing (TGS) (Huang *et al*., 2016), we first performed a comparative sRNA-seq analysis between non-infected (mock inoculated) and infected plants with two Tobamoviruses (TMV-Cg and ORMV) at 4 dpi (Zavallo *et al*., 2015) to evaluate and select genes mapped by 24-nt endogenous sRNAs.

### Selection of a subset of genes with differential accumulation of sRNAs

Original data were fully reanalyzed by means of the new tools. We used Shortstack software (Axtell, 2013) to select genomic regions with differentially accumulated (DA) sRNAs of 24-nt between treatments. This reanalysis allows an improved method of placement of multimapping reads based on weighted probabilities by the unique mapped reads (Johnson *et al*., 2016) (see more on Material and Method section). Then we identified features of interest (genes, promoters or nearby transposons) associated with DA sRNAs. A 1.5 fold-change difference between treatments and a p-value below 0.001 were considered for the selection. Table 1 details the selected genes with 24-nt sRNAs mapped to promoter regions (GP), gene body (GB) or nearby transposons (TEs). Panel a) details differential genes selected by comparing each virus (TMV-Cg/ORMV) with non-infected plants (MI), whereas panel b) shows the comparison between both viruses (ORMV vs TMV-Cg). A total of 211 genes associated to DA sRNA with cluster between treatments with the two analyzed viruses versus the non-infected plants (Table S1). Of these genes, 44% corresponded to genes mapped by sRNAs in gene promoter, 11% in gene body and 45% in genes located near TEs (2kb each side) (Fig. 1a). Additionally, 80 genes showed significant differences between virus (ORMV vs TMV-Cg) treatment. In this case, 14% consisted of genes mapped by sRNAs in gene promoter, 20% in gene body and 66% in TEs near genes (Fig. 1a).

**Figure 1.**
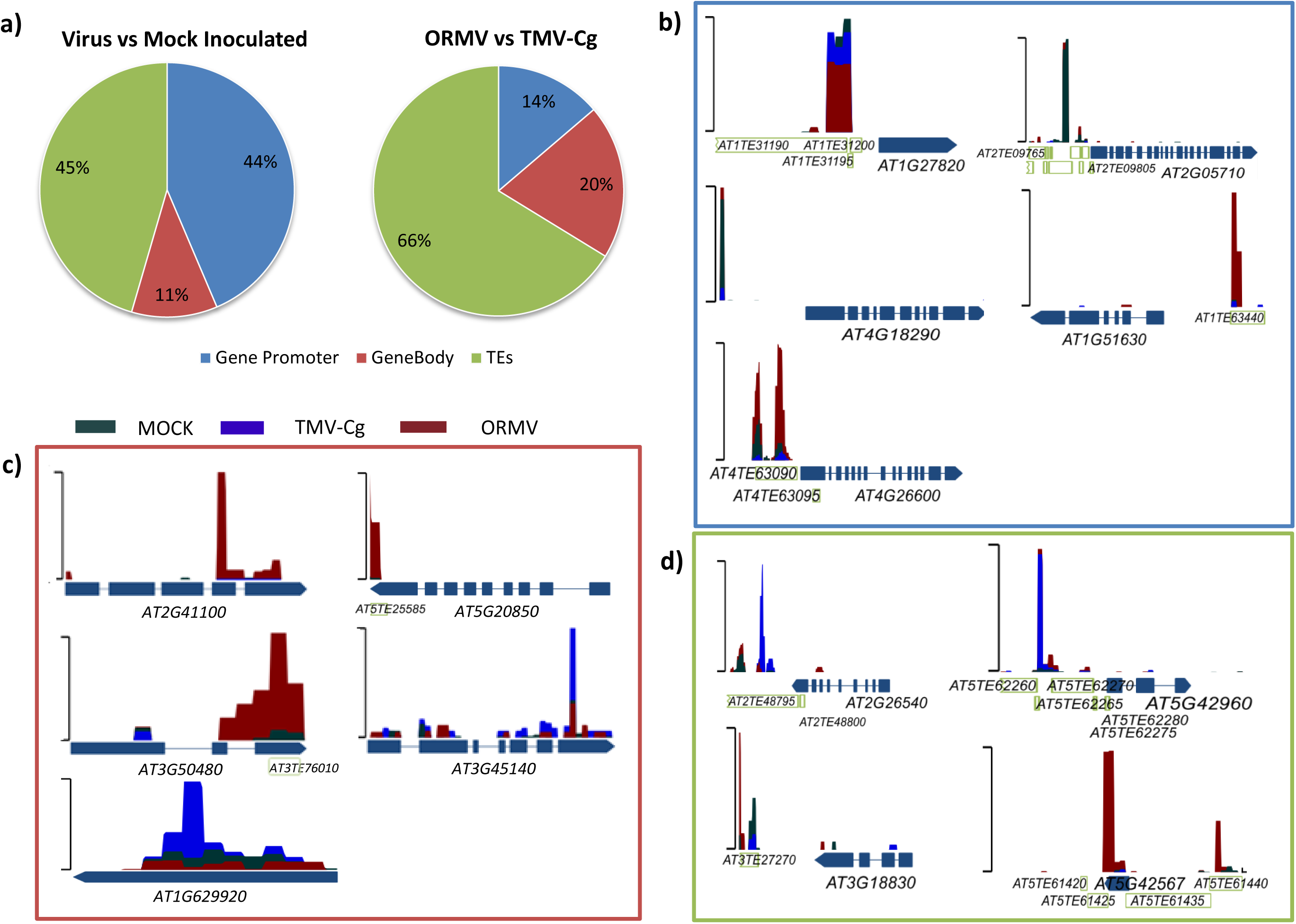
Schematic representation of differential distribution of 24-nt sRNAs in selected genes or different regions in *Arabidopsis thaliana* plants infected with *Tobamoviruses* (ORMV or TMV-Cg). a) Pie graph showing the percentage of differentially represented regions mapped with the analyzed sRNAs: gene promoter, gene body or transposon near genes. Left panel: comparison between infected vs mock-inoculated plants. Right panel: comparison between both strains of viruses. Distribution of 24-nt sRNA mapped to b) gene promoter of the selected genes (blue square), c) gene body (red square) or d) transposons within or nearby genes (green square). The y-axis represents the abundance of sRNAs and the x-axis represents promoter, gene body or transposon regions.

**Table 1.**
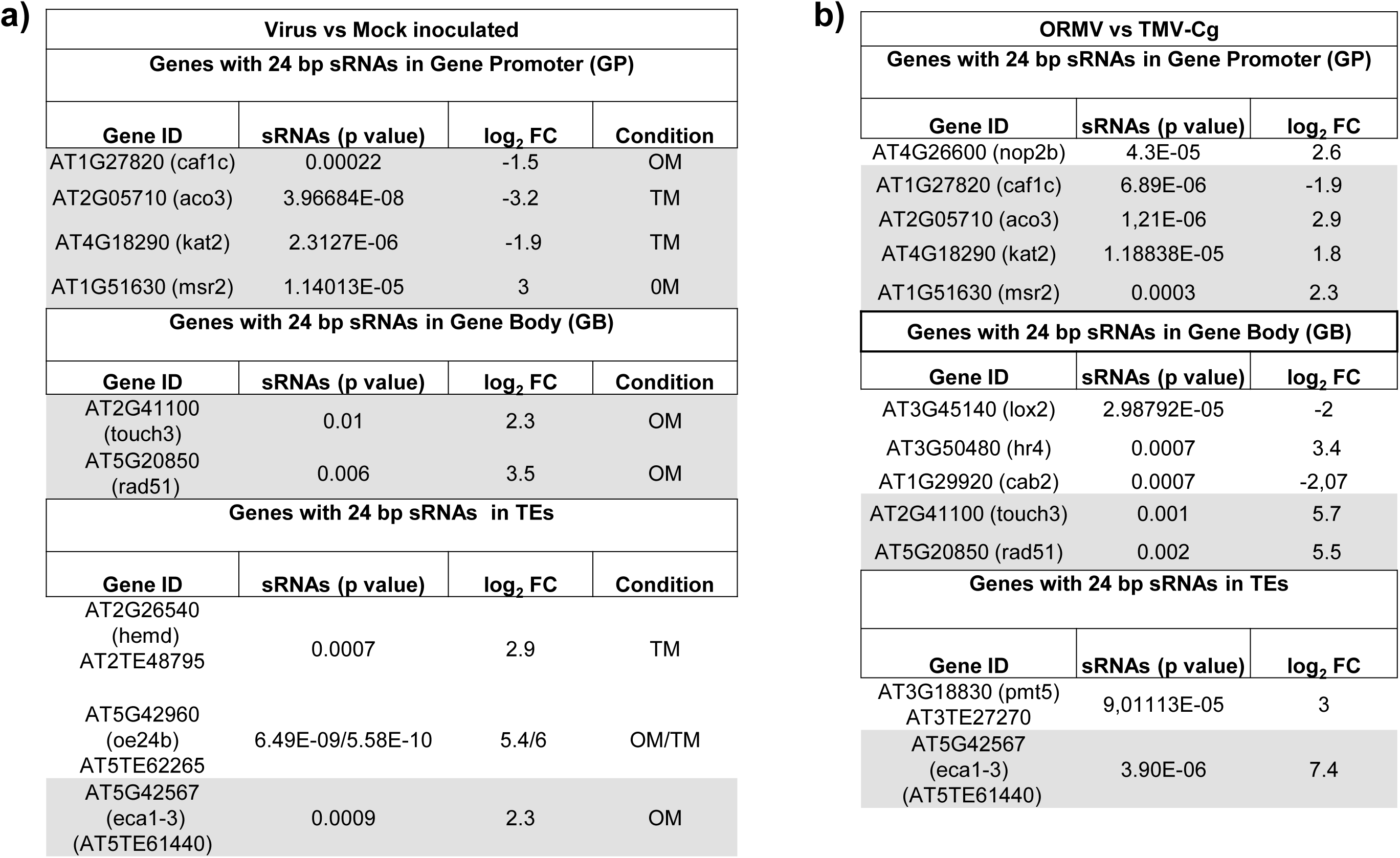
Subgroup of selected genes with differential accumulation of endogenous 24-nt sRNAs in Arabidopsis thaliana infected with two Tobamoviruses (TMV-Cg and ORMV) at 4 dpi. a) sRNAs with differences between ORMV and/or TMV-Cg versus mock-inoculated plants mapped to promoters, gene body or transposons of the selected genes. b) sRNAs with differences between ORMV versus TMV-Cg in *A. thaliana* plants mapped to promoters, gene body or transposons of selected genes. The genes highlighted in gray appear in both tables. M: Mock inoculated; TMV-Cg (T): Tobacco Mosaic Virus; ORMV (0): Oilseed Rape Mosaic Virus.

Figure 1b-d shows the distribution of 24-nt sRNAs in plants infected with TMV-Cg and ORMV and mock-inoculated plants in three different regions of the DNA (set of selected genes of Table 1), either in the promoter (Fig. 1b), in gene body (Fig. 1c) or/and in transposons near (2Kb) non-coding regions (5-UTR and 3-UTR) of chosen genes (Fig. 1d).

In detail, the infection with ORMV and TMV-Cg produced accumulation alteration of 24-nt sRNAs in selected genes. Furthermore, in accordance with Zavallo et al. 2015, we confirmed that viral infection alters the profile of sRNAs in early times of infection. Importantly, viral infection alters to greater extent 24-nt sRNAs associated with TEs (Fig. 1d). We selected four genes close to (2kb) TEs that were differentially mapped by 24-nt sRNAs. Indeed, three of the TEs belonged to the RC/Helitron superfamily: ATREP 3, in the 3-UTR region of the AT2G26540 gene; ATREP1, 2-kb from the 5-UTR region of the AT5G42960 gene and ATREP10D, 1-kb from the 5-UTR region of AT5G42567 gene. Finally, AT9NMU1: DNA/MuDR superfamily located near the 3-UTR region of AT3G18830 gene. It is important to note that most of the selected genes (Table 1) have TEs nearby when considering a region larger than the 2 kb previously selected.

### Molecular analysis of the selected genes in wild type plants

To assess whether the infection with two strains of *Tobamovirus* affects mRNA expression levels of the selected genes, we performed quantitative PCRs, using specific primers for 14 of these genes. At 4dpi most of the selected genes showed increased mRNA levels in the infected plants, except AT3G45140 (lox2), which showed no significant differences (Fig. 2). Moreover, in wild type plants (Col-0), all genes mapped by 24-nt sRNAs within the gene promoter region showed higher levels of mRNA (Fig. 2a), because of the viral infection. Importantly, a negative correlation was evident between sRNA accumulation and transcription levels in most of this group of genes (i.e. a decrease of sRNA amount and an increase of mRNA levels in the infected plants).

**Figure 2.**
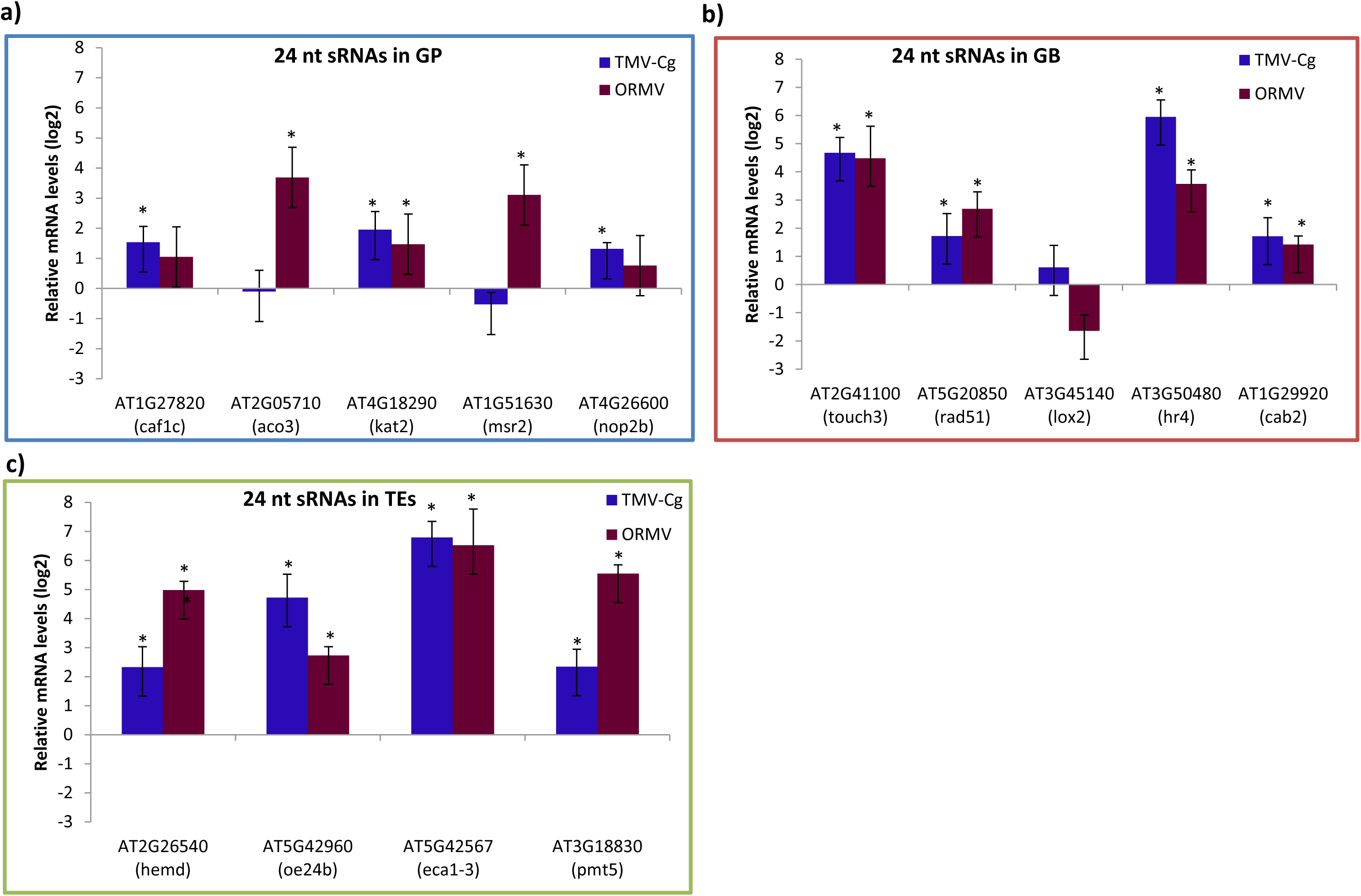
Molecular analysis of target genes in wild type plants (Col-0). qRT-PCR analysis of mRNAs of selected genes at 4dpi of TMV-Cg and ORMV-infected plants (in relation to mock-inoculated plants) with differential accumulation of 24-nt sRNA in a) gene promoter (GP) (blue square), b) in gene body (GB) (red square) or in transposons (TEs) (green square). Log_2_-fold change of selected genes is shown. Error bars represent SE. * = p ≤ 0.05. The blue and red bars refer to plants infected with TMV-Cg or ORMV, respectively.

On the other hand, genes with 24-nt sRNAs mapping to the gene body region exhibited higher relative levels of mRNA and only one these genes showed no differences of expression with any of the evaluated *Tobamoviruses* (Fig. 2b). This latter case can be explained by the antagonism between the SA and JA pathways: lox2 is a JA-biosynthesis gene sensitive to suppression by SA accumulation (Leon-Reyes *et al*., 2010). Interestingly, the expression level of this group of genes showed a positive correlation with sRNA abundance in infected versus non-inoculated plants. Finally, genes with sRNAs mapping to nearby transposons also showed higher relative levels of mRNA in infected plants (Fig. 2c).

These results suggest that viral infection modifies the transcription of different genes and that this could be as a consequence of changes in the profiles of 24-nt sRNAs that mapped to different gene regions or to TEs nearby genes.

### RNA-directed DNA methylation and demethylation mechanism have a role in the transcriptional changes induced by the infection

To assess if the changes in mRNA levels of the selected genes depend on alterations of methylation levels linked to RdDM pathway, we evaluated the expression of these genes in different mutant plants. For this purpose, we used plants with alterations in various components of the RdDM pathway and a mutant in a triple methyltranferase (demethylases). At 4 dpi, most of the selected genes presented an unaltered expression profile after the infection in the three evaluated mutants of the RdDM machinery (Fig. 3). The evaluated mutants were the RNA dependent RNA polymerase 2 mutants, unable to synthesize double stranded RNA: *rdr2.5*, (Fig. 3a); the argonaute 4 mutant, unable to guide methylation processes: *ago4.1* (Fig. 3b); the mutant without a common subunit of polymerases PolIV and PolV: *nrpd2a* (Fig.3c); and a triple demethylase mutant ros1-dml2-dml3, denominated *rdd* (Le *et al*., 2014) (Fig. 3d).

**Figure 3.**
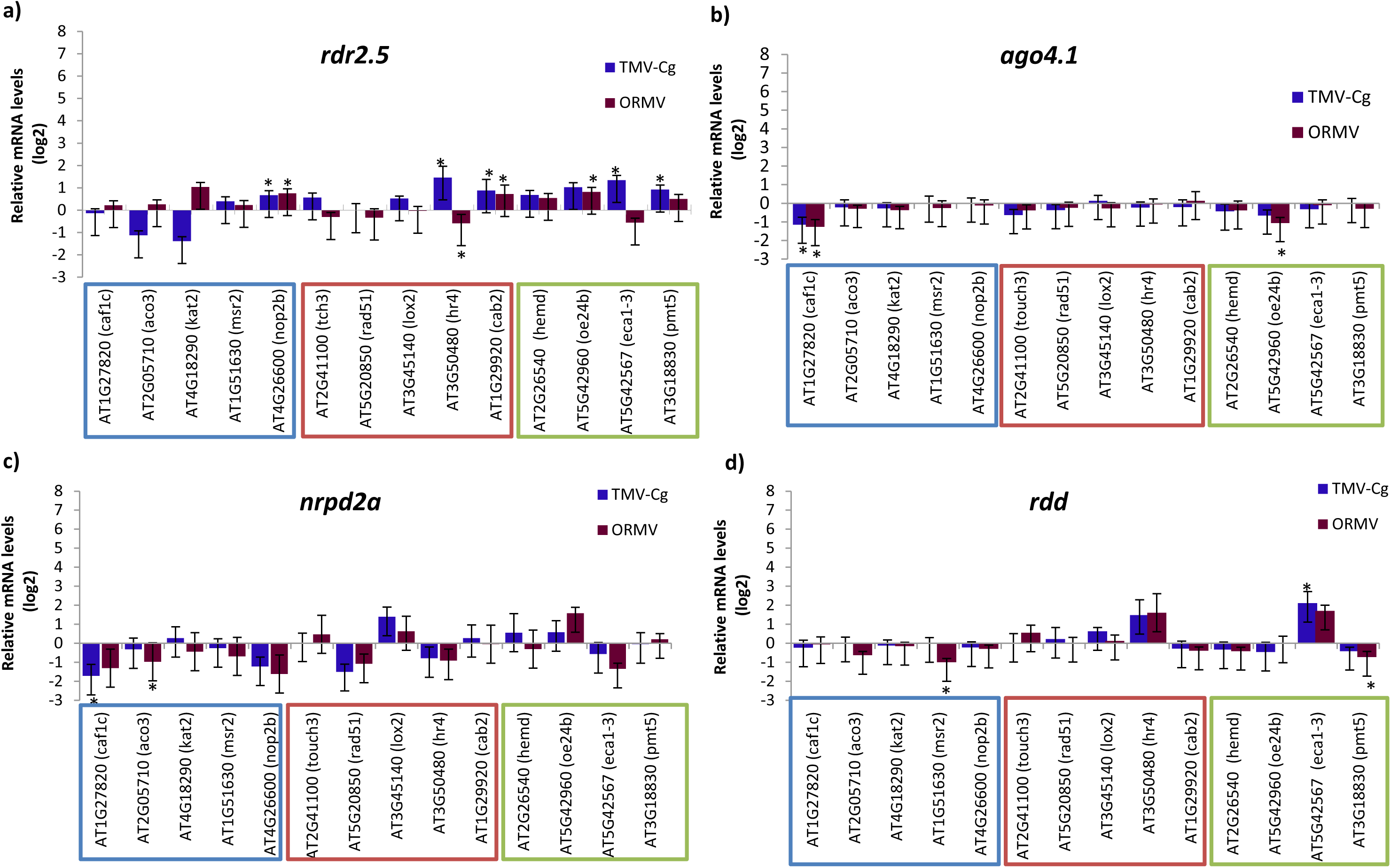
Molecular analysis of target genes of RNA-directed DNA Methylation (RdDM) machinery and triple demethylase mutants. RT-qPCR analysis of selected genes at 4dpi of TMV-Cg and ORMV infected plants in comparison to mock-inoculated plants. Log_2_-fold change of selected genes at 4dpi are shown in a) RNA–dependent RNA polymerase 2 (rdr2.5) mutant; b) argonaute 4 (ago4.1) mutant; c) nrpd2a (subunit of polymerase IV and V) mutant and d) triple demethylase (rdd: ros; dml2; dml3) mutant (. The blue, red and green boxes display the genes mapped by sRNAs in gene promoter, body regions or in transposons, respectively. Error bars represent SE. * = p ≤ 0.05. The blue and red bars show plants infected with TMV-Cg or ORMV, respectively.

Altogether, mRNA levels of the selected genes lost the regulatory impact produced by viral infection because of the disruption of the RdDM machinery or of different demethylases. Thus, the alteration of mRNA levels of the selected genes during viral infection could be regulated by changes in DNA methylation patterns, which in turn are directed by sRNAs.

### Phenotypic evaluation of RdDM and *rdd Arabidopsis* mutants infected with two strains of *Tobamoviruses*

Next, we analyzed the symptoms produced during ORMV and TMV-Cg infection in *rdr2.5*, *ago4.1*, *nrpd2a* and *rdd A. thaliana* plants. The two *Tobamoviruses* exhibited differences in symptom severity in Col-0 ecotype and ORMV produced the most severely altered phenotypes (Zavallo *et al*., 2015).

First, we performed a visual characterization of the symptoms of infected (ORMV or TMV-Cg) and mock-inoculated (controls) plants (Fig. 4). Typical symptoms caused by viral infection are reduced rosette growth, delayed flowering, leaf chlorosis and senescense induction. Col-0 plants inoculated with TMV-Cg showed a smaller rossette size and greater chlorosis than controls and this effect was more pronunced in ORMV inoculated plants.

**Figure 4.**
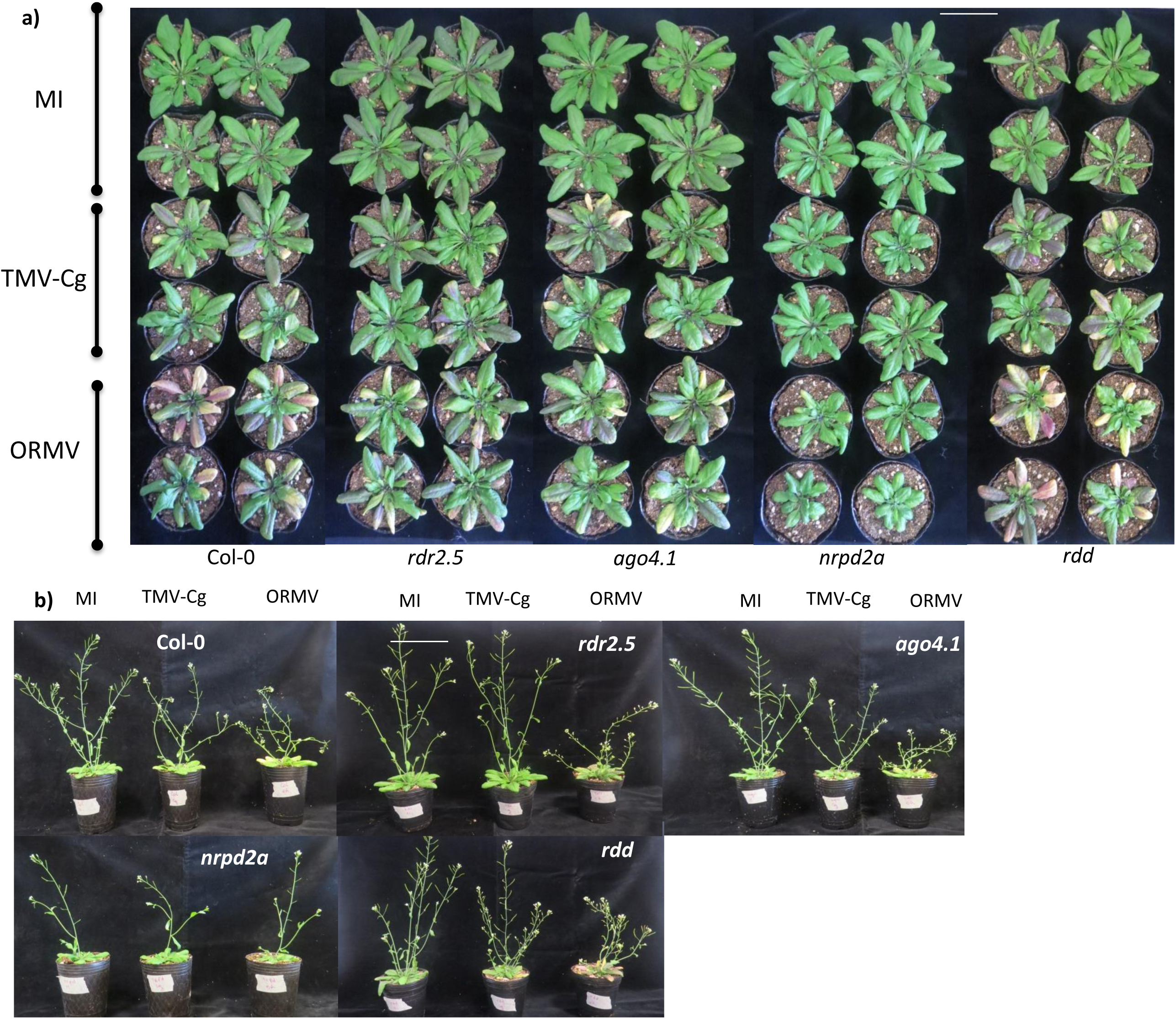
Visual phenotypic characterization. a) Photography of *Arabidopsis thaliana* Col-0 (wild type), RdDM and *rdd* mutants infected with ORMV or TMV-Cg viruses. At 16 days post-inoculation (dpi), the bolts were cut-off the plants before photographing to obtain better images. b) Photography of bolt at 16dpi form plants inoculated with both evaluated viruses. *rdr2.5*: RNA–dependent RNA polymerase 2 mutants; *ago4.1*: argonaute 4 mutants; *nrpd2a*: subunit of polymerase IV and V mutants; *rdd*: triple demethylase (ros; dml2; dml3) mutant. Bar, 1 cm.

Interestingly, *rdr2.5* and *ago4.1* mutants were less affected by the viral infection. At 16 dpi, these mutants showed minor leaf chlorosis, decreased rosette growth inhibition. Furthermore, the severity of the symptoms between the two viruses was similar and less evident than in wild type plants. Particularly, the *nrpd2*a mutant showed no leaf chlorosis, whereas, the triple mutant *rdd* did display this symptom, although with limited growth inhibition (Fig. 4a). Another typical symptom of viral infection is the delayed emergence of bolting. Col-0 ecotype plants inoculated with ORMV showed shorter bolting than TMV-Cg-inoculated plants; similarly, the *ago4.1* mutants presented a comparable disease phenotype than that of wild type plants (Fig. 4a). In addition, the *rdr2.5* and *rdd* mutants showed no delay in bolt emergence when infected with TMV-Cg and, surprisingly, the double *nrpd2a* mutants showed null variation with either of the viruses (Fig. 4b).

In parallel, we performed RT-qPCR assays using specific primers for replicase to assess gene virus accumulation at 7 dpi. Similar ammounts of viral RNA was evident in Col-0 and the evaluated mutants (Fig. 5). Thus, the alteration of the detected symptoms cannot be explained by a diferential viral accumulation. To obtain a more thoroughly phenotypic characterization of physiological impacts of the infection, we measured and quantitated several important traits: bolt height (Fig. 6a), rosette fresh weight (Fig. 6b), chloropyll acumulation (Fig. 6c), anthocyanin acumulation (Fig. 6d), phenolic acumulation (Fig. 6e) and carotenoid acumulation (Fig. 6f).

**Figure 5.**
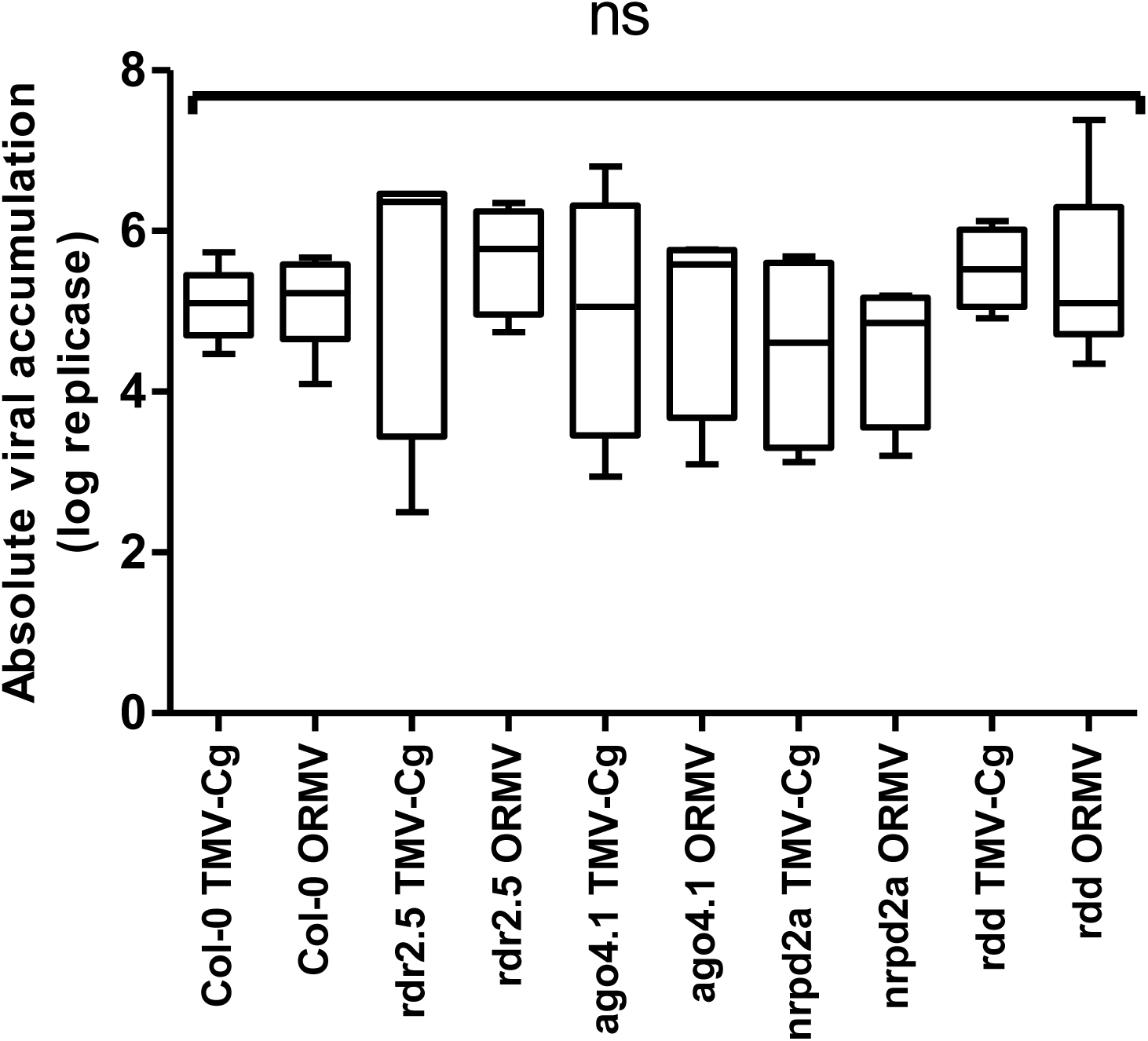
Viral accumulation level. Absolute viral loads for Col-0 and *Arabidopsis thaliana* RdDM/*rdd* mutants inoculated at 7 dpi obtained using *Tobamovirus* replicase specific qPCR primers. ns: no significant differences.

**Figure 6.**
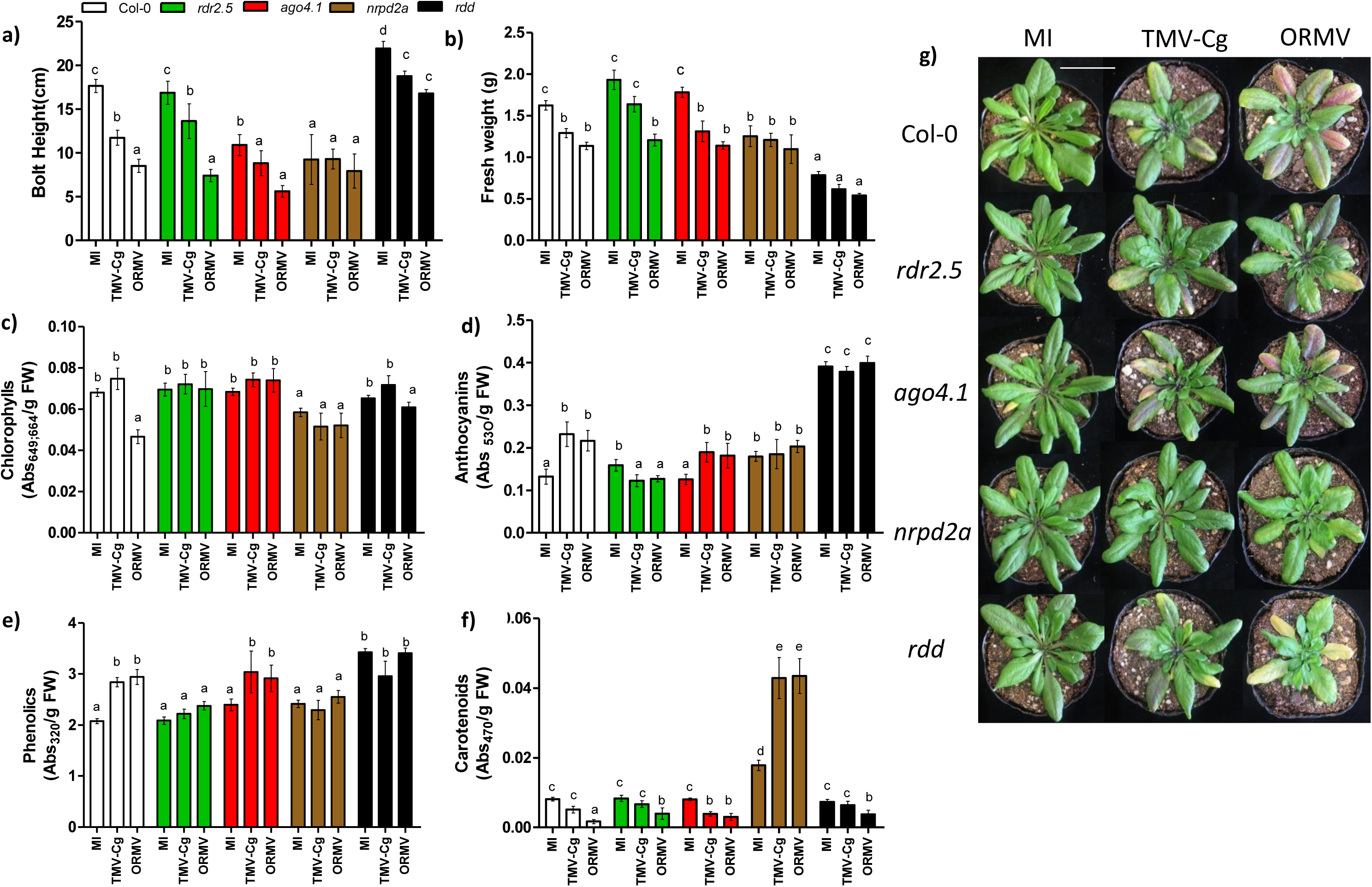
Phenotypic measurements in wild type, RdDM and *rdd* mutants infected with two *Tobamoviruses*. *Arabidopsis thaliana* leaves of the different genotypes were harvested 16 dpi. a) Bolt height; b) Rosette Fresh weight; c) Chlorophyll accumulation d) Anthocyanin accumulations; e) Phenolic accumulation; f) Carotenoid accumulation. The different letters indicate significant differences between MI/TMV-Cg and ORMV inoculated samples. Error bars represent standard error. p≤0.05. g) Photograph of Col-0, RdDM and *rdd* mutants inoculated with both viruses at 16 dpi (the bolt was cut before photographing to obtain better images). Bar, 1 cm.

Relative loss of fresh weight is a marker of senescence that is a common physiological process induced by viral infection. As expected, Col-0 and in *rdr2.5* and *ago4.1* plants inoculated with both *Tobamoviruses* (ORMV and TMV-Cg) presented significant differences regarding fresh weight, althought growth inhibition was lower in both mutants. Interestingly, the *nrpd2a* mutant and the triple mutant *rdd* showed no significant differences in rossette weight in comparison to mock-inuculated plants (Fig. 6g).

Decrease of chlorophyll level in leaves during senescence process is another well-characterized pattern occuring under biotic or abiotic stresses (Khanna-Chopra, 2012; Pružinská *et al*., 2005). To confirm the findings of the visual characterization, we also measured chlorophyll levels of the different mutants infected with the two viruses. No significant differences were detetcted in wild type plants inoculated with TMV-Cg; ORMV infection, by contrast, considerably reduced the amount of chlorophyll at 16dpi. Notably, none of the RdDM pathway mutants showed clear symptoms of senescence except for the methyltransferase mutants, which were affected only by ORMV (Fig. 6c,g).

Anthocyanins are polyphenolic compounds that are induced during senescence under stress and that participate in many processes, including the defense against different pathogens,(Liu *et al*., 2018; Landi *et al*., 2015). Phenolic compounds are secondary metabolites produced by the plant against different pathogens and, like anthocyanins, are produced under different stresses (Mazid *et al*., 2011). Wild type and *ago4.1* mutant plants infected with both Tobamovirus displayed increased levels of anthocyanins and total phenolic compounds. No evident differential accumulation of these compounds, however, was detectable in the other mutants used in this assay (Fig. 6d,e).

On the other hand, carotenoids are antioxidant molecules induced by oxidative stress and their accumulation is reduced during viral infection (Havaux, 2014; Ibdah *et al*., 2014). Col-0, *rdr2.5* and *rdd* plants showed significant differences of this compund only when infected with ORMV. However, the *ago4.1* mutants showed the lowest accumulation of this compound upon infection of any the viruses. Notably, the polymerase IV/V impaired mutant showed increased levels of carotenoids with the two viruses (Fig. 6f).

Altogether, the production of symptoms during viral infection is partly caused as a result of the action of RdDM pathway guided by sRNAs. Problably, this is due to changes that modify the dynamic balance between methylation and demethylation events.

## Discussion

Although the mechanisms by which a viral infection induces symptoms have been widely studied (Dardick, 2007; Jay *et al*., 2011; Inaba *et al*., 2011; Shimura *et al*., 2011; Smith *et al*., 2011; Yang *et al*., 2019; Bilgin *et al*., 2003), the role of epigenetics in the production of symptoms is still unclear. DNA methylation is the major epigenetic hallmark associated to the control of TEs or repetitive elements by means of transcriptional gene silencing, which is mainly directed by 24-nt sRNAs. Here, we tested the effect of 24-nt sRNAs after a viral infection, by emphasizing the role of RdDM pathway in the regulation of symptom induction.

First, viral infection altered the profile of 24-nt sRNAs at early times after infection. The selection of this particular type of siRNAs was due to their main role as guides of transcriptional gene silencing. In accordance with similar studies published in the last years (Guo *et al*., 2017; Zavallo *et al*., 2015), here the viral inoculation modified the balance of 24-nt sRNAs profiles that mapped to coding regions or to transposons located near genes. One possible explanation for this alteration is that, by means of endogenous machinery, the plant defense generates vsRNAs and therefore controls the infection; this defense process simultaneously could affect the levels of other sRNAs. In addition, viral counter-defense actions, like the activation/expression/induction of suppressors of silencing proteins, could be interacting with components of the RdDM pathway, thus affecting their function. For example, Cucumber mosaic virus suppressor 2b interacts with AGO4 protein and reduces its activity (Hamera et al., 2012). Moreover, the RNA decay mechanism and sRNA function have a clear association and, consequently, an alteration of RNA decay impacts in the accumulation of many endogenous genes regulated by silencing (Martínez de Alba *et al*., 2015). Furthermore, Conti et al., (2017b) demonstrated that viral infection alters RNA decay machinery, which in turn leads to impact on the silencing pathways. Therefore, altering sRNA-mediated functionality could be another way to explain the viral impact on the profile of 24-nt accumulation (Gabriela Conti *et al*., 2017).

The analysis of the detected DA sRNA clusters in the infected plants, in comparison to non-infected plants, demonstrated that most of the DA sRNAs mapped to transposons close to genes. To a lesser extent, sRNAs mapped within promoter regions that sometimes contains a TE and, finally, a smaller percentage mapped within gene body regions. We also observed that the abundance of the 24-nt sRNAs that mapped to promoter regions was lower in infected plants in comparison to non-infected plants and that the transcription level of the genes mapped by these sRNAs was higher in Col-0 ecotype.This finding is consistent with the knowledge that DNA methylation in regulatory regions inhibits transcription; therefore, a negative correlation between the amount of sRNAs (associated to a reduced level of methylation) and the level of mRNA is expected (Mette *et al*., 2000; Přibylová *et al*., 2019).

On the other hand, controversy surrounds the effect of methylation of gene body regions. Whereas some researchers reported that the methylation of these regions promotes the expression of certain genes, others demonstrated otherwise (Li *et al*., 2012; Yang *et al*., 2015). Herein, genes with increased abundance of sRNAs in gene body regions showed also higher relative levels of mRNAs in infected plants. These patterns could be explained by a mechanism, proposed by Harris *et al*. (2018), in which the methylation reader SUVH1/3 DNAJ complex enhances transcription.

Furthermore, we identified a common positive correlation between the amount of 24-nt sRNAs and the expression of genes located near TEs. Of the four TEs near genes selected in this study, three belong to Helitron family and the other to the DNA/MuDR family. Notably, almost every evaluated gene (13/14) have TEs near non-coding regions (5-UTR and 3-UTR). Most of the detected TEs correspond to Helitron family. Indeed, researchers have identified both types of TEs associated with genetic and epigenetic variation located in proximity to disease resistance NBS-LRR genes (Kawakatsu *et al*., 2016; Underwood *et al*., 2017; Cambiagno *et al*., 2018).

Altogether, *Tobamovirus* viral infection altered the profile of 24-nt sRNAs and, in turn, these small molecules can regulate gene expression.

**Second**, we confirmed that (de) methylation is an important controller of gene transcription. Indeed, the different knockout mutants in RdDM pathway as well as the *rdd* triple mutant were insensitive to any induction of gene expression because of the viral infection. The suppression of genes required by the methylation machinery guided by 24-nt sRNAs in order to promote TGS inhibited the increase of mRNA levels. By contrast, the wild type plants did present upregulated levels of mRNAs upon *Tobamovirus* infection.

In the first instance, we attributed the differences of gene expression to a lower susceptibility of the mutants to the virus. This hypothesis, however, was discarded because the (de) methylation mutants and wild type plants accumulated similar levels of viruses. Thus, this deregulation may be due to modifications in the RdDM pathway and, therefore, the genes may be directly affected by RdDM machinery or may be indirectly altered by the action of other targets that modify closed and open chromatin states regulated by TGS (Harris *et al*., 2018).

Zhong et al. (2013) demonstrated that the presence of a TE is an important factor for Pol V stably assembles to a given genomic locus. Therefore, Pol V may be associated to regions where promoters and transposons overlap near PolV targets. In addition, the findings in their study indicate that genes close to methylated transposons are often associated with reduced genetic expression (Zhong et al., 2013). In this context, Gagliardi et al. (2019) showed that a MITE like-TE with inverted repeats (IR), located near the WRKY6 locus, dynamically regulates the expression of these genes by altering the chromatin topology. They evidenced a non-canonical mechanism of regulation in which the enrichment of DNA methylation in flanking regions of WRKY6 is associated with the formation of chromatin loops that ends up altering transcription rates. Although a reduction of methylation in these regions together with stable methylation in one intron correlates with the formation of an intragenic loop that inhibites the transcription of the gene (Gagliardi et al., 2019).

Interestingly, most of the genes evaluated in this study had TEs near 5-UTR/3-UTR and 24-nt sRNAs that map to promoter or coding regions. Possibly, the RdDM pathway directs changes in methylation patterns and these changes produce modifications in DNA topology. These variations of topology therefore would open or close regulatory regions, thus promoting or inhibiting the entry of RNA polymerase II and therefore modifying the transcription of these genes.

**Third**, we provide clear evidence that (de) methylation impact on the production of symptoms induced by viral infection. All tested mutants showed differences in at least one of the phenotypic variables measured in comparison with *Tobamovirus*-inoculated Col-0 ecotype, at later stages of infection. Although the *rdr2* and *ago4* mutants are the mutants that most closely resemble Col-0 plants, both had reduced leaf chlorosis. In addition, the *rdr2* mutants also accumulated lower amounts of anthocyanin and phenolic compounds. Plants with mutation in the common NRPD subunit of polymerases IV and V, which are involved in RdDM machinery, showed fewer symptoms (no bolt growth inhibition, no chlorosis and no decrease in fresh weight or in accumulation of anthocyanin and phenolic compounds) in comparison to wild type plants, but strikingly showed very high levels of carotenoids. The triple demethylase *rdd* mutants showed an intermediate phenotype with viral infection: lower fresh weight inhibition and, like Col-0 plants, lower levels of chlorophyll and carotenoids.

If we analyze the RdDM pathway in detail, we can hypothesize that the evaluated *rdr2* and *ago4* mutants produce less symptoms than *nrpd* plants upon viral infection because of the existence of other proteins with functional redundancies or because of the action of the non-canonical pathway of methylation. Indeed, some sRNAs of the PTGS via RDR6 and probably AGO6 could be used as guides to activate RdDM (Cuerda-Gil and Slotkin, 2016). Both polymerases IV and V are major components of the RdDM machinery. Pol IV transcribes sRNAs from target loci and physically interacts with RDR2, whereas Pol V recruits AGO4 and directs the methylation of complementary sequences (López *et al*., 2011; Haag *et al*., 2012). The *nrpd* mutants affect an essential subunit shared by both Pol IV and Pol V and this disruption may explain the almost complete absence of symptoms. Therefore, the mutation may not only lead to the inhibition of methylation, but also to the absence of chromatin remodeling factors that interact with Pol V and regulate the transcriptome.

Furthermore, changes in symptomatology could be due to a global effect of the disruption of the methylation machinery that impairs endogenous TGS regulation. For example, Aconitase (aco) coding gene produces an enzyme that can catalyze the conversion of citrate to isocitrate through a cis-aconitate intermediate. *Arabidopsis thaliana* knockout plants for the aco gene have lower chlorosis and are more tolerant to oxidative stress (Moeder *et al*., 2007). Moeder *et al*., in the same research, demonstrated that *Nicotiana* plants silenced by VIGS (90% reduction in enzyme activity) have reduced pathogenic inductive cell death (Moeder *et al*., 2007). In the present study, all the evaluated mutants displayed lower levels of chlorosis and loss the upregulation of aco3 mRNA levels induced by the infection; which suggests that (de) methylation could regulate this process. The absence of chlorosis in the *nrpd* mutant could be explained by the interruption of two essential components (PoIV and Pol V) of the RdDM pathway. This is a clear difference with other mutants, *ago4* and *rdr2*, with redundancy in protein families (Havecker *et al*., 2010; Willmann *et al*., 2011).

On the other hand, changes in TE methylation located near genes could be affecting DNA topology. A recent study reported that senescence induced by darkness activates young TEs of the DNA and Helitron families (Minerva S. Trejo-Arellano *et al*., 2019). This physiological process is associated with negative histone regulation and replication remodelers that could mobilize these TEs. Minerva S. Trejo-Arellano *et al*. ((Minerva S. Trejo-Arellano *et al*., 2019)) also demonstrated that changes in the regulation of the RdDM pathway alter the senescence process.

Altogether, we propose that chlorosis, a characteristic symptom of viral infection, possible as well as other symptoms, may be regulated by alterations in methylation patterns that affect both TGS and the three-dimensional structure of DNA and all these DNA modifications consequently may affect the levels of transcription.

In conclusion, this research represents the initial step in exposing that DNA methylation directed by endogenous sRNAs has a central role in the production of differential symptoms in plants.

## Experimental procedures

### Plant material

Seeds of Arabidopsis Arabidopsis thaliana were stratified at 4°C for 3 days. All plants were grown under standard conditions in controlled environmental chambers, at 22°C and a 16 h white light: 8 h darkness photoperiod (Boyes et al., 2001). The rdr2.5, ago4.1, nrpd2a and rdd lines were all in the Col-0 background (Ariel et al., 2014).

### Viral infection and sample collection

Arabidopsis plants were mechanically inoculated with carborundum into their third true leaf at stage 1.08 (Boyes et al., 2001). The quantitation of TMV-Cg and ORMV inocula were performed by infecting Nicotiana tabacum (NN) plants with serial dilutions of viral extracts. Local lesions (LL) were counted and the inoculum stored at −80°C until infection. Subsequently, 5 μl of ORMV and TMV-Cg inoculum diluted (100 LL per plant) in 20 mM phosphate buffer (pH 7) was added to the leaf and its surface was gently abraded. The mock-inoculated plants were buffer rubbed instead. The systemic leaves 8 and 11 were sampled 4 days post inoculation (dpi) to isolate RNA and 7 dpi to quantify viral accumulation. Six independent plants were used for each treatment. Pigment measurement was performed in individual leaves 16 dpi.

### Bioinformatic analysis

Original data used in Zavallo et al.’s study were reanalyzed (Zavallo et al., 2015). FASTX-Toolkit (version 0.0.13) was used to perform adaptor and quality trimming on sRNA-seq experiment reads for each library. Bash command was used to filter and select 24nt-length reads, whereas ShortStack software (versión 3.8.5) (Johnson et al., 2016) was used to map and quantitate sRNA reads to the reference Arabidopsis genome TAIR10 assembly. In more detail, cleaned 24nt-lenght reads were mapped to reference genome without the – locifile option. Then, bam files of each treatment were merged with samtools merge into a single bam sRNA mapping file. This file was used to re-run ShortStack using the -bamfile option with different feature annotation files with the –locifile option such as genes, promoters and TEs. Count files of each run were subjected to further analysis to detect differential accumulation (DA) sRNAs between treatments with edgeR package using the glm model. Three sets of DA sRNAs were analyzed: DA sRNAs within genes, DA sRNAs within promoters (minus 2.5KB from gene transcription start site) and DA sRNAs within TEs (located up to 2 kb of the 5-UTR and 3-UTR regions of the selected genes). Software deepTools (version 3.3.1) was used to perform sRNA distribution plots of selected features.

### RNA isolation and Real time quantitative polymerase chain reaction (qPCR)

Total RNA was isolated from frozen Arabidopsis thaliana leaf tissues using TransZol Reagent (TransGen Biotech, Beijing, China) and treated with DNAse I (Invitrogen, California, USA). Subsequently, cDNA was synthesized using MMLV (Invitrogen, California, USA) and random primers according to the manufacturer’s instructions (Invitrogen). All qPCR experiments were carried out in an ABI StepOne Plus Real Time PCR System (Applied Biosystems, California, USA). Table SI1 displays the oligonucleotide primer sets used for qPCR. Arabidopsis EF1alpha gene was used as reference for normalization in each experiment. For each gene measurement, six biological replicates were used for each gene measurement in each treatment. qPCR data analysis and primer efficiencies were obtained using LinReg PCR software (Ruijter et al., 2009). Relative expression ratios and statistical analysis were performed using fgStatistics software interface, which uses the 2-ΔΔCt method to analyze the relative changes in gene expression (Di Rienzo, 2009; Livak and Schmittgen, 2001). A p value below 0.05 was set as the cut-off for statistical significance.

### Physiological and pigment measurements

First, bolt height of the different treated plants was measured with a ruler, whereas fresh weight was determined with an analytical balance. Pigments were extracted from individual samples of leaf 5 at 16 dpi. Chlorophylls and carotenoids were extracted using 96% ethanol. The measurement of anthocyanins and soluble phenolic compounds was performed by placing the already weighed leaves in acidified methanol (99: 1, v/v) at 4°C for 48 h (Rabino and Mancinelli, 1986; Leone et al., 2014). Absorbance of 0.2 ml was measured at 664, 649, and 470nm (chlorophylls and carotenoids), 530 and 657 nm (anthocyanins) or 320 nm (phenolic compounds) in a Multiskan Spectrum (Thermo Fisher Corp., Waltham, MA, U.S.A.). Chlorophyll concentrations were calculated using the following equations: Ca+b = total chlorophylls (µg/ml) =1496 / 22.24 × A649 + 5.24 × A664, Ca = chlorophyll a (µg/ml) = 13.36 × A664 – 5.19 × A649, and Cb = chlorophyll b (µg/ml) = 27.43 × A649 – 8.12 × A664. Carotenoid concentrations were assessed using Cc = total carotenoids (µg/ml) = (1,000 × A470 – 2.13 × Ca – 97.64 × Ca)/209. Anthocyanin contents were evaluated using A530 – 0.25 × A657. The experiment was repeated three times independently.

### Statistical analyses

Statistical analyses were performed with one-way ANOVA test using INFOSTAT software (InfoStat/Professional version 2008, Grupo InfoStat. FCA, Universidad Nacional de Córdoba, Córdoba, Argentina, https://www.infostat.com.ar/). The significance level for all tests was α = 0.05. Appropriate transformations of the primary data were used when needed to meet the assumptions of the analysis. The phenotypic characterization was assessed by analyzing the data by factorial ANOVA, with viral treatment and genotype as factors.

## Acknowledgements

This research was supported by grants from CONICET (Consejo Nacional de Investigaciones Científicas y Técnicas) and ANPCyT (Agencia Nacional de Promoción Científica y Tecnológica) PICT 2014-1163 and PICT 2015-1532. We thank Federico Ariel, Pablo Manavella and Martin Crespi for the *rdr2*, *nrpd2a*, *ago4* and *rdd* lines, respectively.

## Author contributions

ML: Investigation, Methodology, Visualization, Writing original draft and Writing review and editing. DZ: Investigation, Bioinformatics analysis, Data curation, Writing review and editing. AV: Investigation, Methodology, Writing review and editing. SA: Conceptualization, Funding acquisition, Investigation, Project administration, Resources, Supervision, Writing review & editing.

## Short legends for Supporting Information

Table S1. Normalized reads of 24-nt sRNAs in mock-inoculated (MI), TMV-Cg and ORMV infected plants at 4dpi. Each excel sheet shows the different readings that map to the different regions. Cpm: counts per million.

Table S2. Oligonucleotide primers used for RTqPCR experiments.

